# Interpreting neural decoding models using grouped model reliance

**DOI:** 10.1101/656975

**Authors:** Simon Valentin, Maximilian Harkotte, Tzvetan Popov

## Abstract

The application of machine learning algorithms for decoding psychological constructs based on neural data is becoming increasingly popular. However, there is a need for methods that allow to interpret trained decoding models, as a step towards bridging the gap between theory-driven cognitive neuroscience and data-driven decoding approaches. The present study demonstrates *grouped model reliance* as a model-agnostic permutation-based approach to this problem. Grouped model reliance indicates the extent to which a trained model relies on conceptually related groups of variables, such as frequency bands or regions of interest in electroencephalographic (EEG) data. As a case study to demonstrate the method, random forest and support vector machine models were trained on within-participant single-trial EEG data from a Sternberg working memory task. Participants were asked to memorize a sequence of digits (0–9), varying randomly in length between one, four and seven digits, where EEG recordings for working memory load estimation were taken from a 3-second retention interval. Present results confirm previous findings in so far, as both random forest and support vector machine models relied on alpha-band activity in most subjects. However, as revealed by further analyses, patterns in frequency and particularly topography varied considerably between individuals, pointing to more pronounced inter-individual differences than reported previously.

**Author summary:** Modern machine learning algorithms currently receive considerable attention for their predictive power in neural decoding applications. However, there is a need for methods that make such predictive models interpretable. In the present work, we address the problem of assessing which aspects of the input data a trained model relies upon to make predictions. We demonstrate the use of grouped model-reliance as a generally applicable method for interpreting neural decoding models. Illustrating the method on a case study, we employed an experimental design in which a comparably small number of participants (10) completed a large number of trials (972) over multiple electroencephalography (EEG) recording sessions from a Sternberg working memory task. Trained decoding models consistently relied on alpha frequency activity, which is in line with existing research on the relationship between neural oscillations and working memory. However, our analyses also indicate large inter-individual variability with respect to the relation between activity patterns and working memory load in frequency and topography. Taken together, we argue that grouped model reliance provides a useful tool to better understand the workings of (sometimes otherwise black-box) decoding models.

## Introduction

The application of statistical algorithms to neural data is becoming an increasingly popular tool for explaining the link between biology and psychology [1, 2]. Supervised learning algorithms, in particular methods such as random forest (RF) [3] and support vector machines (SVM) [4] algorithms, are frequently utilized to decode various psychological phenomena related to, for instance, perception, attention, and memory with promising success [5–9]. However, while these algorithms are optimized to provide accurate predictions, their interpretability is often not given.

While encoding models aim to model the brain’s response to stimuli, decoding models can be used to efficiently assess the presence of decodable information in a certain brain area [10]. In the simple case a of linear models and under some assumptions, a transformation of the weights allows a decoding model to be interpreted as an encoding model [11]. As complex models cannot be interpreted as directly, however, there is a need for methods that allow researchers to understand what drives these model’s predictions [9, 10, 12]. One application where such interpretations are required is in the case of exploratory data-driven analyses, to gain an insight into which parts of the data ^1^ an accurate model uses to guide a closer examination of these relationships. Furthermore, having developed a predictive model, researchers may be interested in assessing the plausibility of a trained model in relation to existing empirical research and theoretical work, or for troubleshooting unexpected predictions. It should be noted that care has to be taken when trying to interpret decoding models as “reading the code of the brain”, as decoding alone does not provide a computational account of information processing in the brain [10].

The present approach focuses on model-agnostic interpretations, targeting the question of which parts of the data a trained decoding model relies upon to make predictions. Here, model-agnosticism refers to an interpretation that is independent on the particular class of models being used [13]. For instance, random forest and SVM models are based on different principles (ensembles of decision trees for random forests and optimally separating hyperplanes for SVM). Rather than interpreting models in terms of their parameters, which may not be easily comparable in the case of different model classes, a model-agnostic method allows to interpret the influence of a predictor variable in any supervised model.

Usually, the importance of predictors in a multivariate model is assessed for individual predictors, such as partial regression coefficients in a linear regression model, Gini importance or permutation importance in random forest algorithms [3]. However, as variables extracted from EEG or MEG recordings, for instance, are often inter-correlated, questions about the importance of predictor variables rather concern sets of conceptually related variables than individual variables [14, 15]. For instance, when assessing the reliance on certain topographical or spectral components for predicting a psychological phenomenon, neural activity may be shared across multiple brain regions or recording sites. The acquisition resolution of these components is usually on a more detailed level that is used for interpretation [16]. Hence, a method that assesses the reliance on groups (or subsets) of variables provides researchers with more meaningful “chunks” for interpretation [13].

A practical approach for assessing the reliance on variables is given by permutation importance, used initially as a measure for the importance of variables in the Random Forest algorithm [3]. The reliance on a variable is quantified by the decrease in prediction performance when that predictor variable is randomly permuted, essentially “nulling” the association between that predictor variable and the outcome. An intuitive terminology for this idea for any learning algorithm is given by *model reliance*, as proposed by Fisher et al. [17]. Model reliance indicates the extent to which a model relies on specific variables in making predictions, i.e. the extent to which performance decreases when permuting that predictor variable. Crucially for the present problem, the method of permuting predictor variables can be adapted to permuting groups of conceptually (or statistically) related variables (such as frequency bands, as opposed to single frequencies) to measure their aggregate impact on predictive performance, as proposed by Gregorutti et al. [18]. This is required as generally, the reliance on a group of variables is not equivalent to the sum of individual model reliances [18]. To emphasize that the interpretation of a variable’s (or group of variables’) influence in a model’s prediction is based on the particular model being used, the term model reliance is adopted in this work, following Fisher et al. [17]. By design, this approach treats the model as a black-box, thereby making it a model agnostic method that can be used for any supervised learning algorithm.

In order to demonstrate the use of grouped model reliance on a well-established construct in cognitive neuroscience, random forest and SVM models are employed in this work to decode working memory load based on single-trial electroencephalographic (EEG) data, collected in multiple experimental sessions per participant. Consequently, grouped model reliance is used to interpret models in terms of conceptually meaningful groups of features from a single-subject perspective.

Working memory is a widely studied psychological construct and refers to the temporary retention of information in the absence of sensory input, needed for a subsequent behavioral outcome. Neuronal oscillations are hypothesized to be involved in working memory by generating a temporal structure for the brain [19, 20]. Amplitude modulation of neuronal oscillations, in particular in the alpha frequency bands (8-14Hz), is a robust finding in psychophysiological research [21–23]. It is hypothesized that these power modulations aid the functional brain architecture during retention, protect against interference and thereby manifest in relevant behavioral outcomes, as measured by accuracy and reaction time [24, 25]. Thus, the scaling of alpha power with working memory load is considered as an essential neural manifestation of the psychological construct of working memory. Apart from alpha activity, most prominently oscillatory activity in the theta (4-7Hz) and gamma (60-80Hz) frequency bands have been linked to working memory. It has been proposed that theta-band oscillations underlie the organization of sequentially ordered working memory items, whereas gamma-band oscillations are thought to contribute to the maintenance of working memory information [24, 26–30].

Although oscillatory activity from different frequency bands have been established as correlates of working memory across individuals, some studies suggest that inter-subject variability may be high. This variability, however, can take different forms. For instance, working memory load-dependent shifts in alpha peak frequency have been shown to vary between individuals with low versus high working memory capacity [31]. There is also evidence for individual differences in the exact frequency range in which the alpha-rhythm is modulated during the exertion of working memory [32]. In comparison, for theta activity, power modulations have been reported to vary substantially between subjects [33–35] as well as between trials of individual subjects [36]. There is no consensus, however, on the determinants of this inter-subject variability. As a way forward, employing single trial EEG analysis as well as assessing decoding models on a single-subject level may be able to provide complementary information to that of group-level statistics [37–40].

In the present study, the Sternberg working memory task is used [41, 42]. Compared to other paradigms, this task has the advantage that the periods of encoding, retaining and recognizing stimuli are all temporally separated [24, 25]. Subjects are first presented with a list of items, the number of which determines the working memory load. Following a retention interval of several seconds, a probe item is presented, and subjects indicate the membership of this item to the previously presented list.

In the present study, using single-trial EEG data from a Sternberg task, random forest and SVM models are trained on individual subjects to perform working memory load estimation based on power spectra from the retention period. Consequently, grouped model reliance is used to interpret the trained models. In order to put the interpretations of decoding models into the context of more traditional methods from cognitive neuroscience, cluster-based statistics are employed to further probe the relationship between working memory and neural oscillations.

## Materials and methods

### Participants

Eleven subjects were recruited by advertisement at the University of Konstanz (*M* = 24.5 years, *SD* = 4.8; 50% female) and reported no history of neurological and/or psychiatric disorders. One subject was excluded from the analysis, as data acquisition was interrupted during the second session of the experiment. All participants gave written informed consent in accordance with the Declaration of Helsinki prior to participation. The study was approved by the local ethics committee.

### Stimulus material and procedure

Participants performed a Sternberg task [41] with alternating levels of difficulty (1, 4, or 7 items to be kept in memory) while the electroencephalogram (EEG) was recorded. The data of each participant were collected on three different sessions, which were, on average, *M* = 4.35 days (*SD* = 2.72) apart. Written informed consent was obtained from each subject prior to each session. One session comprised four practice trials and six main blocks, each consisting of 54 trials (lasting approximately nine minutes). In between blocks, participants were allowed to rest for a maximum of three minutes. Each participant completed 324 trials per session, resulting in 972 trials in total. Participants were asked to memorize a sequence of digits (0–9), varying randomly in length between one, four or seven different digits. After an initial central fixation interval of 500 ms, the sequence of digits was presented serially. Each digit was presented for 1200 ms, followed by a blank screen for 1000 ms before the presentation of the next digit. After a 3000 ms retention interval (blank screen), a probe stimulus was presented in the center of the screen for 5000 ms. Participants were instructed to indicate whether the probe was part of the previously presented sequence. The right arrow key on a standard keyboard indicated ‘yes’ and the left arrow key ‘no’ answer. Participants’ response was followed by positive or negative feedback for 500 ms. Finally, a blank screen was presented for 1000 ms, after which the next trial began. Within each block, there were nine positive trials (probe part of the study list) and nine negative trials (probe not in the sequence) for each sequence length. Trials were presented in random order with respect to the sequence length.

### Data acquisition

EEG was recorded with an ANT Neuro 128-electrode system (www.ant-neuro.com) with Ag/AgCl electrodes placed on a Waveguard cap with an equidistant hexagonal layout. Signals were sampled at 512 Hz, and electrode impedance was kept below 20 kOhm. The recording was DC and referenced to a common average reference.

### Preprocessing

Data analysis was performed with the MATLAB based FieldTrip toolbox [43]. For each participant and channel, after demeaning and removing the linear trend across the session, independent component analysis (ICA) [44] was used to remove variance associated with eye blink and cardiac activity. Increased noise in the electrodes closest to the ears (LM, LE1, LE2, RM, RE1, RE2) in some participants led to the exclusion of these electrodes from all subsequent analyses for all participants. All trials per session and condition were included. Spectral analyses were conducted for each trial using a Fast Fourier Transformation (FFT) with a single Hanning taper for the retention interval of 3 sec. The predictor variables used for the classification model covered the frequency bands (delta 1–4Hz, theta 5–7Hz, alpha 8–12Hz, beta 13–20Hz and low gamma 21–40Hz) in one Hz steps, for electrodes from nine regions of interest (ROI). Hence, there were a total of 4880 predictor variables (40 frequencies for each of 122 electrodes). ROI’s were left/central/right occipital, left/central/right central and left/central/right frontal. The exact electrodes per group and a layout of their respective locations can be found in Fig 1.

**Fig 1.**
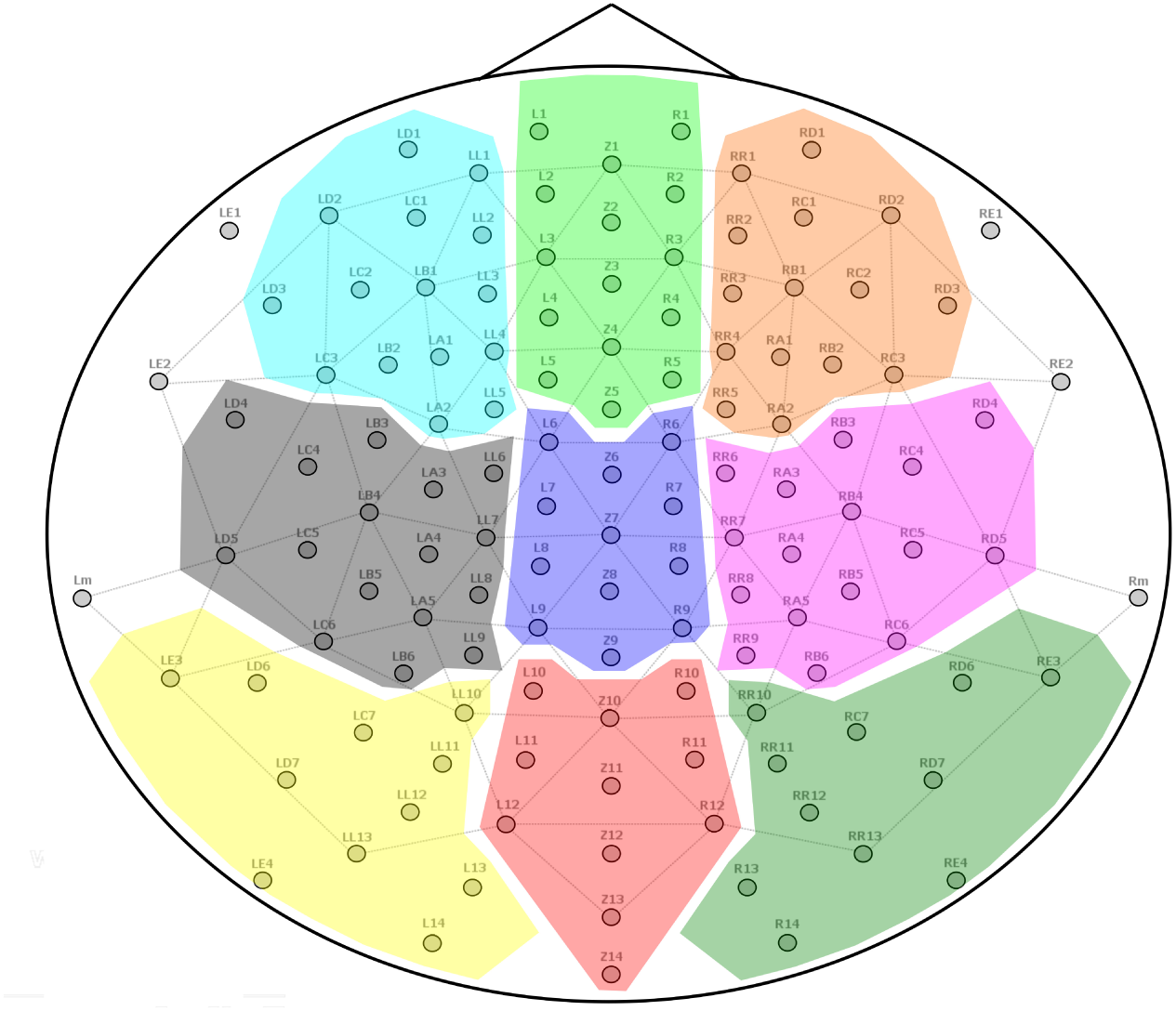
Definition of ROI in terms of electrodes. Note that electrodes LM, LE1, LE2, RM, RE1 and RE2 were excluded from the analyses.

### Decoding model training and evaluation

The Random Forest algorithm, a type of ensemble method, was used as the main model for all decoding analyses [3]. This algorithm was chosen for its ability to perform multiclass classification on a large number of possibly correlated and non-linearly associated variables [3]. The number of trees in the forest was set to 5000 with all other hyperparameters set to default values. Additional analyses employed an SVM model [4] with a radial basis function (RBF) kernel with the penalty term *C* set to 1. Classification accuracy was used as the performance metric for all models. Hence, no distinction was made between misclassifying a load 1 trial as load 4 or 7, for instance.

As the classification task comprises three balanced classes (load 1, 4 and 7), chance level accuracy corresponds to 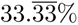. Models for each subject were trained and tested using stratified and shuffled 10-fold cross-validation. Stratification ensures that the distribution of classes is the same for each fold of the cross-validation and can lead to more stable performance estimates than standard *k*-fold cross validation [45]. Simulation studies indicate the most robust performance for stratified *k*-fold cross-validation with *k* set to *k* = 10 [45], which is therefore employed in the present analyses. The reported accuracy and model reliance values correspond to the arithmetic means over all 10 folds of the cross-validation loop. Single-trial data from all 3 sessions and 6 blocks per session were pooled for each participant and used in the cross-validation procedure.

In addition to the within-subject decoding models, between-subject analyses were carried out using a random forest model. Here, models were trained in a 10-fold cross-validation procedure, where the splits were given by individual participants. That is, each training fold consists of all trials of all participants but one, whose trials provide the validation fold. All decoding analyses were implemented in Python, making use of the scikit-learn [46] module.

### Model reliance

Model reliance scores for any particular predictor variable are here defined as a ratio of the error obtained using a random permutation of that variable and the error using the original predictor variables [17]. Note that it is also possible to define MR as the difference in original and permuted error [17]. However, since decodability can considerably differ between participants, the ratio was chosen here for comparability. As such, higher positive MR value for a predictor variable indicates that the model relies on that variables more strongly to make predictions, whereas values towards zero indicate that performance is not impacted by “nulling” the information contained in that variable. Negative MR values can arise due to the randomness of performing a random permutation, but substantial negative values indicate that performance rather improves when the information contained in that variable is permuted randomly [17]. In the present study, the interpretation of model reliance outlined above still holds, but is generalised to groups of variables, rather than individual variables.

Grouped model reliance is then normalized in order to make differently sized groups of variables comparable [18]. This follows the rationale that a large group of variables (such as the gamma-band in the present study) is penalised for its size relative to a smaller group of variables (such as the alpha-band). To this end, the MR score for a particular group of variables is divided by the number of variables in that group.

In the present study, MR is computed on the validation folds in a 10-fold cross-validation procedure. It should be noted that MR could also be computed on the training folds, in which case the interpretation would relate to which variables the model relies upon to fit the training data. This, however, that would depart from the focus of the present study to assess which variables a trained model relies upon to make predictions.

More formally, as adapted from [18], *X* is a *n* by *p* matrix of observations of predictor variables, respectively. **y** is a vector of outcomes of length *n*. *f* is a fitted model. A group of variables (that is, columns) in *X* is indexed by a set *J*, where all *j ∈ J* are 1 *≤ j ≤ p*. ACC_bl_ refers to the baseline accuracy of *f* on *X* and **y**, whereas ACC_perm_*_J_* refers to the accuracy on the data after randomly permuting all predictor variables within each column indexed by *J*. Note that while classification accuracy is used in the present study, MR can be computed on other performance metrics in classification or regression settings. The model reliance value, MR(*X_J_*), for a group of variables, *J*, is given by

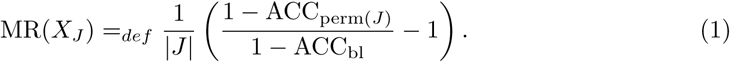

For every cross-validation fold, grouped model reliance is averaged over 10 random permutations and subsequently averaged over all 10 cross-validation folds. This follows from two considerations: While computing grouped model reliance over all possible random permutations is computationally prohibitive, computing MR from only one permutation may lead to unreliable results. As a simple Monte Carlo estimate, averaging over a number of random permutations thus provides a feasible middle ground. Note that this only involves permuting the data from a given validation fold and predicting the class labels rather than re-training the model. Averaging MR over cross-validation folds further provides an estimate of the expected reliances from (partially) different training folds, different validation-folds and random initializations. Thus, the MR scores reported here can be seen as an estimate of the expected reliance for a particular model class, where a class is given e.g. by Random Forest models or SVM) on a particular set of observations. It should be noted that only average MR scores are used here.

Following related work [47], one may also look at computing confidence intervals or *p*-values using the null-distribution of model performance on permuted features. While the interpretations of average MR on the present data did not differ between using 10 or 100 random permutations, one may need more random permutations to obtain reliable estimate of p-values or measures of dispersion. Since obtaining estimates of the variance from cross-validation folds is problematic [48], this may be most relevant for cases where there is a sufficient number of observations to perform a training(-validation)-test split rather than k-fold cross-validation.

Code for implementations of grouped model reliance in addition to Jupyter notebooks for all analyses are available from https://github.com/simonvalentin/wmdecoding.

### Non-parametric statistical testing with clusters

Effects of working memory load on neural data were probed by a cluster-based randomization approach [49]. This approach identifies clusters (in frequency and space) of activity on the basis of which the null hypothesis can be rejected, while addressing the multiple-comparison problem. The null hypothesis tested here was that the trials per subject sampled from the three load conditions stem from the same distribution; thus the labels (i.e., load 1, load 4 and load 7) are exchangeable. Dependent samples F-tests were used as test statistics. Random permutations of the labels were computed 1000 times resulting in a distribution of 1000 F-values. The original value of the test statistic was compared against this randomization distribution using an alpha level of 5%.

## Results

### Behavioral

In order to establish that the experimental manipulation yielded the expected behavioral effects, accuracy, as well as reaction times (RT) of participants, were recorded. In line with previous findings, both accuracy and reaction times decreased with an increase in working memory load (cf. Table 1).

**Table 1.** Means and standard deviations of reaction times and accuracies across subjects.

| Load | Reaction time [ms] |  | Accuracy [%] |  |
| --- | --- | --- | --- | --- |
|  | *M* | *SD* | *M* | *SD* |
| 1 | 530.9 | 133.8 | 97.6 | 2.9 |
| 4 | 659.8 | 147.8 | 95.3 | 3.2 |
| 7 | 784.1 | 144.6 | 89.9 | 6.7 |

Values per participant were computed as the average across all trials.

### Model reliance

Averaged across all subjects, within-subject classification accuracy using the random forest model was 48.51% (*SE* = 1.25%) in a three-class classification task with a chance level accuracy of 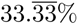. As illustrated in Fig 2 (right) overall, trained models relied mostly on the alpha frequency band. As described in the methods, model reliance is normalized according to the group size of predictor variables. Hence, large groups, such as the gamma band are penalized more than smaller groups, as, e.g. alpha. However, as reported in S1 Fig, even when no normalization for the group size is used, the interpretation that alpha-band activity is central for decoding performance holds. While model reliance for alpha band frequencies consistently emerges across subjects, there is variability in individual profiles, as presented in Fig 3A.

**Fig 2.**
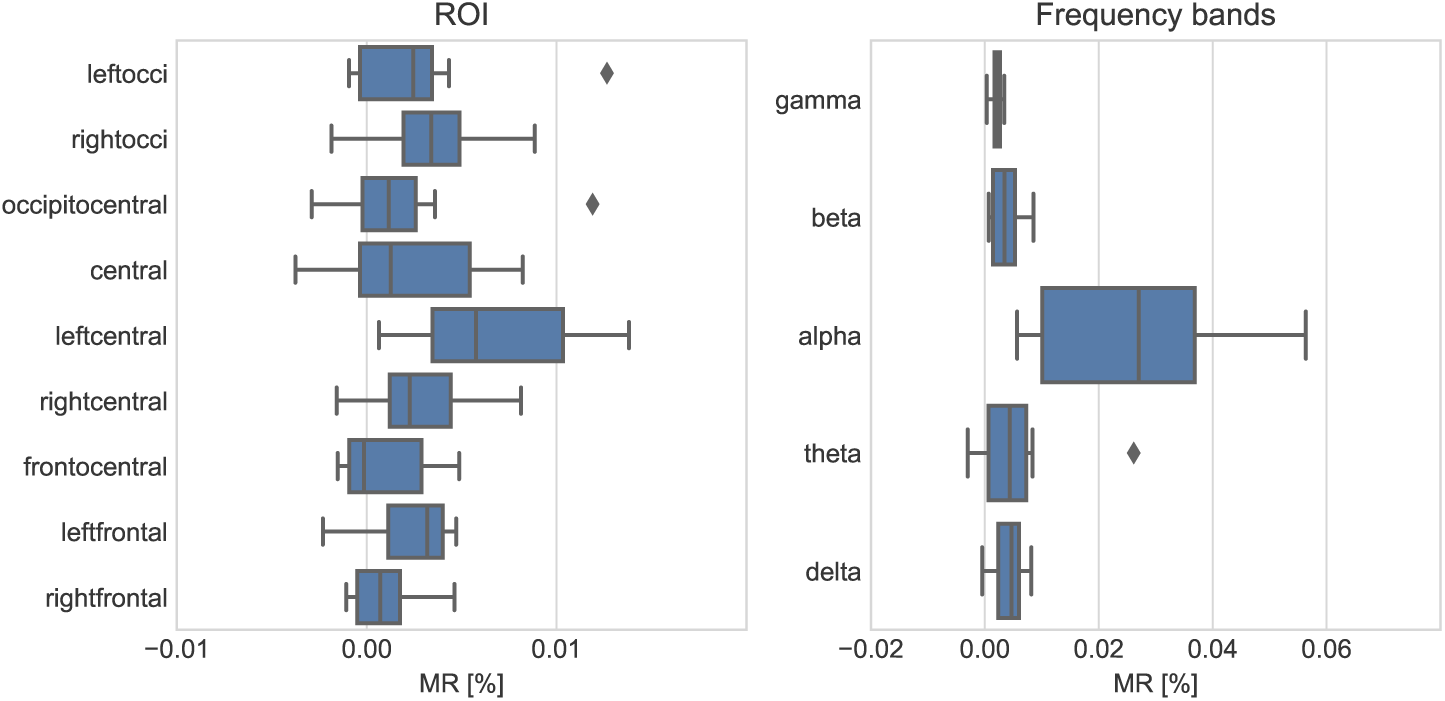
Grouped model reliances. Box-Whisker plots of average grouped model reliance (MR) per participant for different ROI’s (left) and frequency bands (right) using a random forest model.

**Fig 3.**
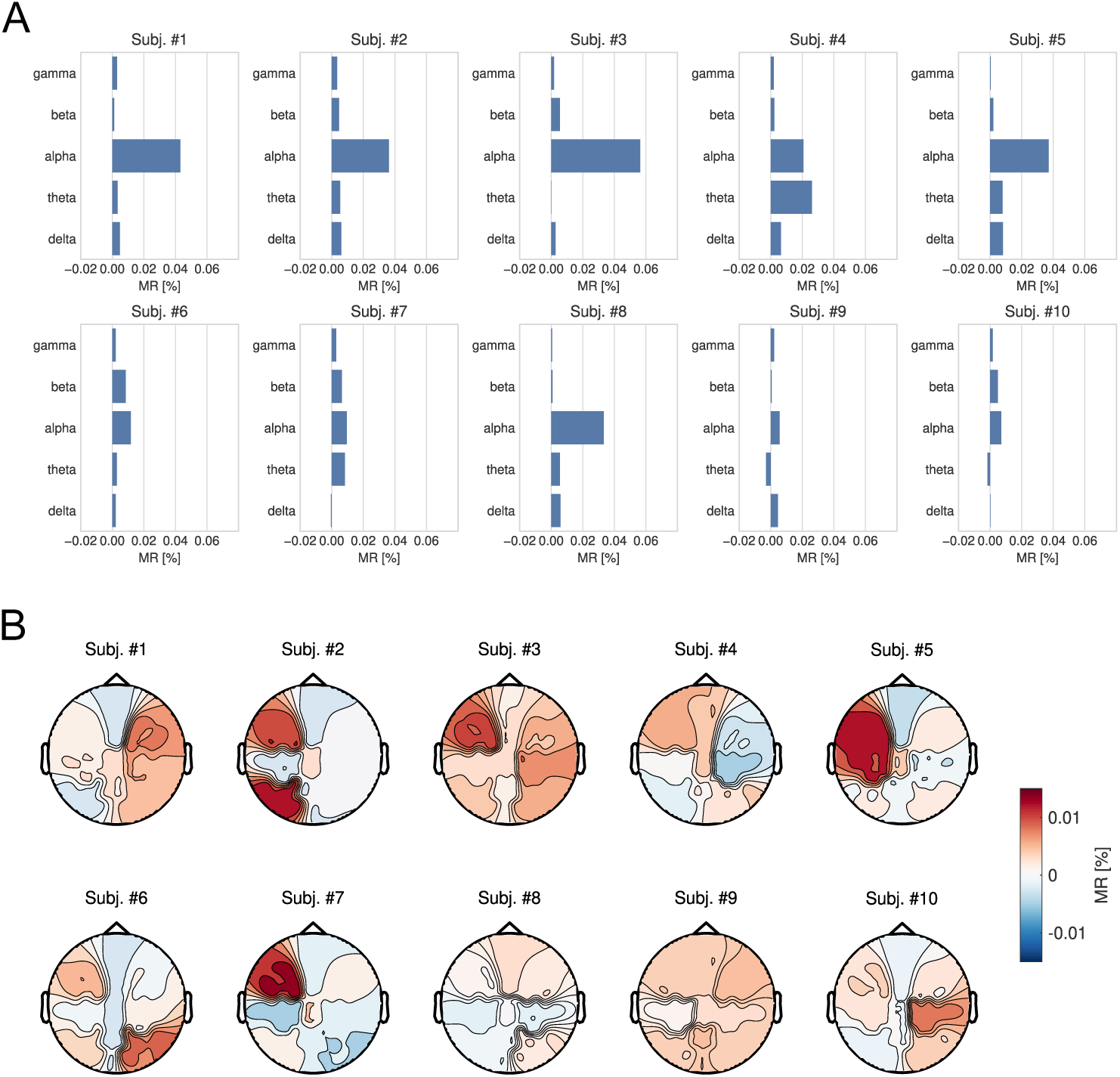
Grouped model reliances per subject. (A) Grouped model reliances (MR) for each frequency band and individual participant. (B) Topography of reliances for each individual participant.

In contrast, MR on scalp topographic features for classification accuracy is less decisive Fig 2 (left). There is no ROI that clearly stands out across subjects in terms of grouped model reliance. Instead, scores show considerable inter-individual variability, as presented in Fig 3B.

To assess whether fitting a different, but similarly performant model would result in comparable estimates of model reliance, additional analyses using an SVM algorithm were conducted. These analyses yielded similar results and interpretations, as illustrated in S3 Fig.

Further, in order to assess whether this reflects only the reliance on these groups of features in a multivariate model, or also the relevance of the feature groups assessed separately, additional analyses were run. Here, a classifier was trained and tested only on separate feature groups (S2 Fig). This analysis further strengthened the interpretation that alpha band activity encodes information that is particularly relevant for decoding working memory load, while model reliance is broadly distributed across ROI.

In support of the notion of individual specificity and intra-individual variation, training and testing a random forest model between subjects yielded comparably poor generalization performance (average accuracy 34.53%).

Additionally, in an exploratory fashion, Spearman rank correlations were computed to assess whether reliance on the alpha-band per participant is associated with performance on the Sternberg task. No statistically significant correlations were found at the 5% level between the reliance on the alpha-band and average reaction time across conditions per subject (*ρ* = *−*0.042; *p* = 0.907), the difference between high and low-load reaction times (*ρ* = *−*0.406; *p* = 0.244), participants’ average accuracy across all conditions (*ρ* = 0.273; *p* = 0.446) or the difference in accuracies between high and low load (*ρ* = 0.37; *p* = 0.293). However, it should be kept in mind that these correlational analyses are based on only 10 data points.

### Cluster-based inferential statistics

Model reliance scores imply that predictors from the alpha frequency group are of particular importance (being the most critical frequency band for all subject except for Subj. #8). Thus, cluster-based statistics were computed on the alpha-band for each participant. As shown in Fig 4A, a significant effect of working memory load on alpha activity was found in all subjects but one (Subj. #9). Descriptively, no clear topographic pattern could be identified across participants.

**Fig 4.**
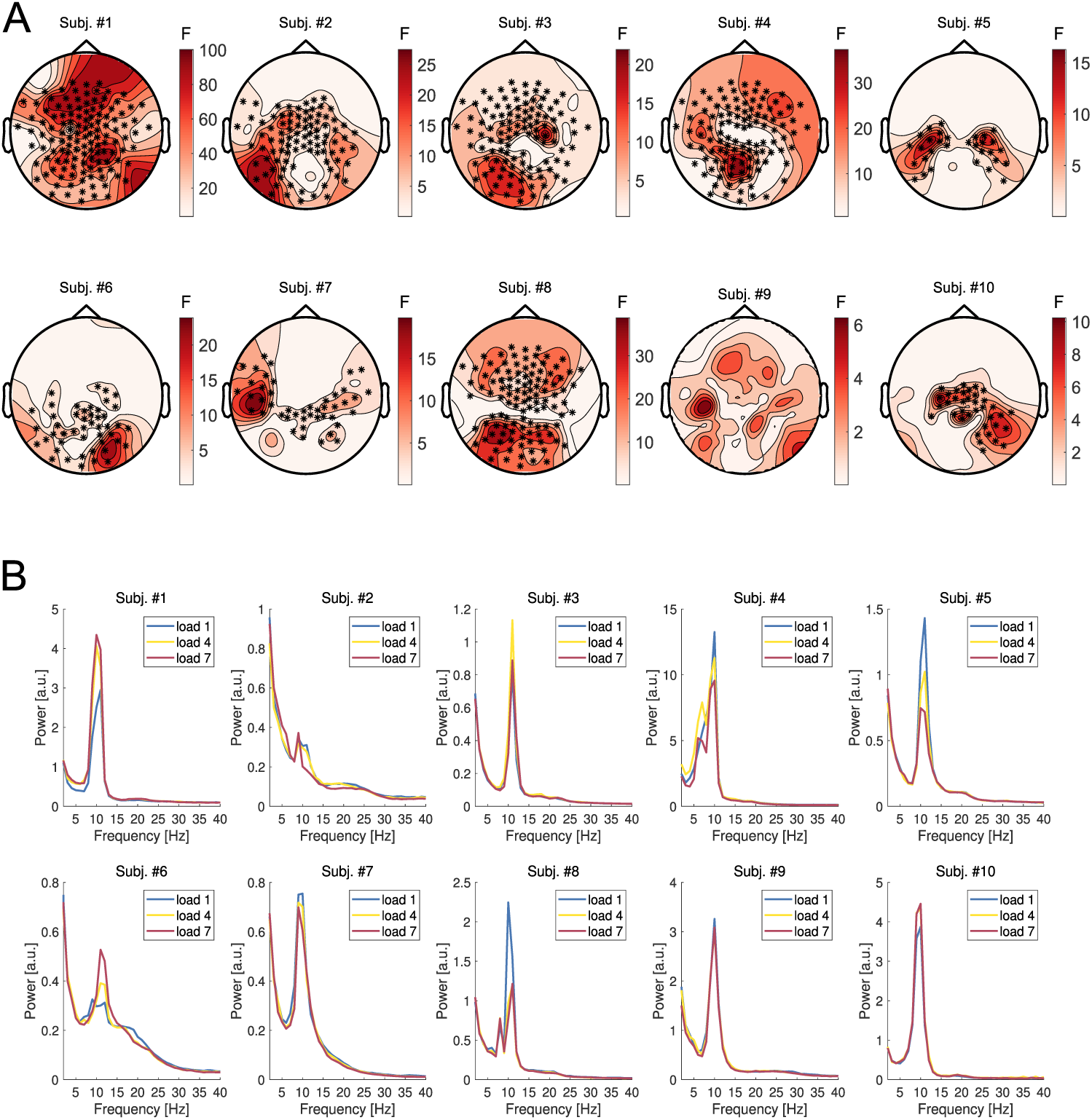
Cluster-based inferential statistics. (A) Topography of the main effect “Load” illustrated for each individual participant. Warm colors indicate the spatial distribution of F-values. Asterisks denote electrodes corresponding to clusters on the basis of which the null hypothesis is rejected. (B) Power spectra averaged across the electrodes belonging to the corresponding clusters illustrated in A. Note that scales are plotted on an individual level, as condition differences within participants are of primary interest.

Power spectra were computed for those electrodes contained in clusters for which significant condition differences were found (Fig 4B)^2^. Crucially, some participants were characterized by a positive relationship of alpha-band activity with increasing working memory load, yet others displayed a reverse ordering or very small to no differences.

Additional analyses were conducted using cluster-based statistics computed across subjects for the alpha and theta frequency bands, for which no significant effects of working memory load were found.

## Discussion

The aim of the present study was to demonstrate the use of grouped model reliance for interpreting decoding models, based on the case study of single-trial EEG recordings from a Sternberg working memory task. Models were probed and interpreted in terms of frequency components as well as ROI on a single-subject level. Decoding models performed with, on average, 48.51% (*SE* = 1.25%) accuracy in a three-way classification task of working memory load. Grouped model reliance scores suggest that across most participants, the alpha band was particularly important for predicting working memory load. Alpha was the most critical frequency band for all participants but one (Subj. #4 for whom theta activity was most important). Further, across participants, models did not rely on particular ROI more than on others. Instead, grouped model reliance scores were found to be distributed across different ROI. To put these interpretations of decoding models into the context of more traditional methods from cognitive neuroscience, subsequent analyses were carried out using cluster-based permutation tests. Here, testing on a single subject level revealed a significant effect of working memory load on alpha power for all but one subject (Subj. #9) However, in contrast to previous accounts, the amplitude of alpha activity increased with load in some individuals (e.g., Subj. #1) while it decreased in others (e.g. Subj. #5). When cluster-based permutation tests were employed on an across subject level, no significant effect of working memory load was found.

Taken together, results from the cluster-based permutation tests are in conflict with previous studies reporting scaling of alpha amplitude with working memory on an across subject level [21, 22, 24, 50]. Instead, the present study identified high inter-individual variability of alpha amplitude and topography. Notably, when decoding models were trained across subjects, generalization performance was comparably poor (accuracy 34.53%), supporting the interpretation of high heterogeneity between subjects. Additional analyses therefore aimed to test whether this observed heterogeneity relates to differences in behavior, as has been proposed previously [51, 52]. Here, it was found that the reliance on the alpha-band did not correlate with average reaction time across conditions per subject, the difference between high and low-load reaction times, average accuracy across all conditions or the difference between accuracies on the high and low load condition. One interpretation for these findings is the variability found in grouped model reliance does not necessarily arise from differences in cognitive abilities, but from differences of the physiological manifestation of working memory, as well as behavioral strategies used by each individual. In line with this, previous work has shown that individuals who are more likely to employ a verbal, rather than a visual, processing approaches exhibit different neural activation during the Sternberg task [53, 54]. However, findings on how differences in working memory performance relate to task-specific strategies are mixed. For instance, it has been reported that subjects who used a verbal rather than a spatial strategy perform better in a 2-back working memory task [55]. In comparison, for a digit span backwards task, which is similar to the Sternberg task used in the present study, no relation was found between the task-specific strategy and working memory performance [53, 54].

Apart from the alpha-band, theta-band power modulations are commonly reported in the study of working memory load [21, 24, 33] and are hypothesized to play a crucial role in organizing sequential information [22, 24]. In the present study, decoding models for most subjects did not rely on theta, with the exception of subject #4. This might be due to a high variability of theta-band activity, which has been reported both between subjects [21, 35], as well as between individual trials [36]. For instance, in contrast to the seminal study by Jensen and Tesche [33], which found theta power to increase with working memory load in the delay period of the Sternberg task, a subsequent study could not replicate this finding [21]. More precisely, although a frontal theta power increase was present in the group average data, this increase was largely driven by only one subject [33, 34]. Indeed, the high inter-subject variability of theta power reactivity has motivated some studies to pre-screen human subjects for the presence of a theta response prior to conducting the main experiment [34, 56]. Hence, the present finding of theta being most critical for the decoding of working memory load in only one out of 10 subjects might be in line with previous reports on the inter-subject variability of theta power modulation. Note that supplemental analysis using cluster-based permutation statistics revealed no statistically significant effect of working memory load on theta power modulations across subjects. From these findings one cannot infer that theta modulations were absent in all subjects in the present study, however. Instead, high inter-trial variability of theta power modulations might result in decoding models relying less on theta but more on other feature groups, i.e. the alpha-band.

Looking at decoding models more generally, while they have become increasingly popular, a number of methodological and interpretational considerations should be kept in mind. First, relevant to the interpretation of grouped model reliance, decoding models that have predictive power should not be directly interpreted as models of the generative process of the data. That is, decoding models, as used in the present study, are primarily useful to indicate that there is information in the data that allows for classification/regression [10]. Grouped model reliance allows to assess which parts (i.e. which variables or groups of variables) of the data a model relies upon. However, note that this interpretation is relative to the model. For instance, a model may not rely upon groups of variables that contain redundant information already contained in other variables. In such cases, we may make false-negative inferences in concluding that a group of variables is not associated with the outcome if its reliance is (close to) zero. To assess this aspect on the present data, models were also trained and validated on separate groups of frequencies and ROI, leading to broadly similar interpretations.

Additionally, care has to be taken with the interpretation of “information is present” that can be obtained from decoding analyses. Crucially, a decoding model may use various kinds of information, which might take a different form than what one may expect from the perspective of cognitive neuroscience. For instance, similar to suppression effects, a decoding model may give different weights depending on the noise covariance structure of the data [11]. This aspect is discussed in-depth by Hebart and Baker [2], who argue that a distinction can be made between an *activation-based* and *information-based* view on neural data analysis. The activation-based view focuses on patterns of de- and increases of activity (e.g., alpha power) and is typically adopted in cognitive neuroscience. The information-based view, on the other hand, is not restricted to activation but examines any change in the multivariate distribution of the data as information that can be used for making predictions, such as the noise distribution [2, 11]. Given that any information contained in the predictor variables may be used by the supervised learning algorithm to make predictions, preprocessing also plays a role in removing known confounding signals from the data. For instance, in the present case-study, ICA was used to remove ocular and cardiac artifacts from the EEG recordings.

Since model reliance provides a summary of the extent to which a model relies upon particular variables to make predictions, this encapsulates both direct associations with the predicted class (or dependent variable more generally) as well as potentially complex interaction terms. This has the advantage of providing a concise summary of the reliance on a group of variables, but has the caveat of not being able to distinguish between different types of information. For instance, it may be that certain variables are only relevant in a potentially highly complex interaction term with other variables, but not on their own. Hence, rather than false-negative inferences from concluding that a variable is not relevant as discussed above, care also has to be taken with interpreting what it means for information to be present.

Some methods such as linear models allow for inferences about a certain type of information more directly [11] but have the downside of being limited in their flexibility to fit relationships that may be present in the data [57]. Grouped MR has the advantage of being model-agnostic, i.e. it is applicable to any supervised model, and can thus be used on models that may make use of complex non-linear relationships. Further methodological development may build on work by Henelius et al. [58], who propose a permutation-based algorithm to identify groups of variables that interact to provide predictions. As proposed by Fisher et al. [17], one may also be interested in conditional MR, that is, the extent to which a model relies upon a particular variable while holding all other variables constant. To this end, only those observations of a variable are permuted that have the same values on all other variables. While this is comparably straight-forward a small number of discrete variables, the problem of matching variables becomes considerably more intricate with more and particularly with continuous variables, though see [17] for directions.

Looking at fitted models more generally, given that there can be multiple similarly performant solutions in high-dimensional data [59], model reliance, and hence interpretations, may also vary across models. In the present study, cross-validation and repeated random permutations were employed to obtain a representative value of what an “average” model relies upon. Fisher et al. [17] further propose model class reliance as a method to obtain upper and lower bounds on the model reliance of a particular variable for all well-performing models of a certain class, such as a Random Forest or SVM. How these or other approaches of assessing the characteristics of the data a model relies upon in more detail may be applied to grouped MR and used on neuroimaging data is beyond the scope of the present article but may be a fruitful direction for future research.

## Supporting information

**S1 Fig. Model reliance without normalization for group sizes.** Results reveal that alpha still emerges as the group of variables that trained models relied upon the most when not accounting for the size of the group of variables.

**S2 Fig. Classification accuracy for training and testing on individual feature groups.** Results highlight the difference between relevance and reliance on predictor variables in multivariate models.

**S3 Fig. Model reliance for SVM model.** Model used an RBF kernel function, with penalty term *C* set to 1. Cross-validated classification accuracy was 48.21%.

## Acknowledgments

We thank Brigitte Rockstroh for comments on the manuscript.

## Footnotes

1 Note that we use the term *predictor variable* or when the context is clear just *variable* rather than *feature* in this work to stay consistent with conventions in cognitive neuroscience rather than machine learning.

2 As no cluster was found for Subj. #9, power spectra were computed over electrodes selected from Subj #7. This individual was chosen due to the similar topographic pattern of the effect of load.

